# Evolution of transmission mode in conditional mutualisms with spatial variation in symbiont quality

**DOI:** 10.1101/315119

**Authors:** Alexandra Brown, Erol Akçay

## Abstract

While some symbioses are always mutualistic or parasitic, others have costs and benefits that depend on environmental factors. The environmental context may itself vary in space, in some cases causing a symbiont to be a mutualist in one location and a parasite in another. Such spatially conditional mutualisms pose a dilemma for hosts, who might evolve (higher or lower) horizontal or vertical transmission to increase their chances of being infected only where the symbiont is beneficial. To determine how transmission in hosts might evolve, we modeled transmission evolution where the symbiont had a spatially conditional effect on either host lifespan or fecundity. We found that over ecological time, symbionts that affected lifespan but not fecundity led to high frequencies of infected hosts in areas where the symbiont was beneficial and low frequencies elsewhere. In response, hosts evolved increased horizontal transmission only when the symbiont affected lifespan. We also modeled transmission evolution in symbionts, which evolved high horizontal and vertical transmission, indicating a possible host-symbiont conflict over transmission mode. Our results suggest an eco-evolutionary feedback where the component of host fitness that a conditionally mutualistic symbiont influences affects its distribution in the population, and, through this, the transmission mode that evolves.

## 1 Introduction

Most, if not all, multicellular organisms live in symbiosis with other species. While some symbioses are always mutualistic or parasitic, many others have costs and benefits that are context-dependent (Chamberlain et al., 2014; Daskin and Alford, 2012; Thomas et al., 2000). We call these interactions conditional mutualisms. Symbiont effects may vary based on abiotic factors (e.g. nutrient availability (Cheplick et al., 1989) or temperature (Baker et al., 2013)) or biotic factors (e.g. the presence of a third species which parasitizes the host (Smith, 1968)). The abiotic or biotic context may in turn vary in space. In some cases, the symbiont may change from a mutualist to a parasite depending on the location. For example,the endophytic fungus *Epichloë coenophiala* increases the biomass of tall fescue (*Festuca arundinacea*) seedlings in nutrient-rich soil, while decreasing host biomass in nutrient-poor soils (Cheplick et al., 1989). Different temperatures produce a similar pattern in the nutrients provided by the *Symbiodinium* endosymbionts of corals. Clade D members of *Symbiodinium* provide less nitrogen than Clade C symbionts except at high temperatures, where they provide equivalent nitrogen and more carbon (this pattern is thought to explain the geographic distribution of Clade C and Clade D symbioses) (Baker et al., 2013).

Such spatially conditional mutualisms pose a dilemma for hosts in deciding how to acquire their symbionts. In general, assuming no correlation between horizontal and vertical transmission, hosts are predicted to evolve reduced vertical (parent-to-offspring) transmission of parasites and increased vertical transmission of mutualists (Yamamura, 1993). Hosts may also evolve decreased susceptibility to horizontal transmission of parasites, including when resistance comes at a cost to fecundity, if reproduction is local (Best et al., 2011). Hosts may even evolve decreased horizontal transmission of parasites to others in a spatially structured population (Débarre et al., 2012). However, hosts in spatially conditional mutualisms have to deal with a symbiont that is both a mutualist and a parasite, and it is not clear whether horizontal transmission, vertical transmission, both, or neither will evolve in conditional mutualisms. Furthermore, symbionts as well as hosts may show genetic variation that affects the two rates of transmission (Ebert, 2013). There may thus be host-symbiont conflict over transmission mode, which may also influence transmission evolution.

Which transmission mode evolves is an important question, since transmission mode itself, regardless of whether it arises through host or symbiont evolution, influences symbiont spread and the evolution of symbiont costs and benefits. Horizontal transmission is predicted to select for more parasitic symbionts, and vertical transmission for more mutualistic ones (Alizon et al., 2009; Ewald, 1987), in the absence of feedbacks selecting for mutualism (Akçay, 2015; Shapiro and Turner, 2014) or parasitism Werren et al. (2008). Furthermore, research on the impact of spatial variation on parasitism shows that spatial heterogeneity can have a large influence on the virulence and spread of parasites (Carlsson-Granér and Thrall, 2015; Gibson et al., 2016; Jousimo et al., 2014; Lively, 2006; Penczykowski et al., 2014; Real and Biek, 2007; Saeki and Sasaki, 2018; Thrall and Burdon, 2000). While studies of spatial heterogeneity in parasitism have generally focused on spatial variation in host or parasite traits, distribution, or transmission (but see (Krist et al., 2004; Tellier and Brown, 2011), which include environmental effects on the costs of infection), they suggest that spatial heterogeneity can have an important impact on symbioses. Understanding transmission mode evolution in hosts and symbionts in spatially conditional mutualisms may thus give insight into both potential host-symbiont conflict as well the future distribution and virulence of the symbiont.

We model transmission mode evolution in a spatially conditional mutualism over a range of newborn host dispersal rates. We consider two different types of spatially conditional mutualisms that affect different components of host fitness. In the first conditional mutualism, the symbiont affects host lifespan, and in the second the symbiont affects host fecundity (modeled as chance of reproduction per unit time). We split symbiont effects into these components partly because they lead to significantly different evolutionary predictions, and partly because it is possible that a symbiont may have a strong effect on one component but not the other. For example, symbioses that are involved only with reproduction, like plant-pollinator/seed parasite relationships will influence host fecundity without affecting lifespan. On the other hand, symbioses involved with, for example, juvenile survival (as in the interaction between jellyfish and the juvenile scads they protect from predators) affect lifespan without having any influence on the reproductive output of hosts who survive to adulthood (Bonaldo et al., 2004). (We also consider several examples of conditional mutualisms affecting host lifespan and fecundity in the supplement.) To determine whether there is host-symbiont conflict over transmission, we model transmission mode evolution under host and symbiont control separately. We infer the possibility of conflict if hosts evolve one transmission rate and symbionts evolve another.

Intuitively, we may predict that when a host is likely to stay in the same location as its parent, vertical transmission may be a good strategy to ensure an advantageous infection status (i.e. infection where the symbiont is beneficial and lack of infection where the symbiont is harmful). Conversely, when hosts often disperse from their natal patch, they might instead rely on horizontal transmission from their new neighbors to acquire the symbiont where it is beneficial. However, horizontal transmission will only confer the “right” infection status when a host’s neighbors are infected where the symbiont is a mutualist and uninfected where the symbiont is a parasite. Thus, hosts should only evolve horizontal transmission when the distribution of infected hosts matches the spatial distribution of symbiont effects. As the distribution of infected hosts is itself influenced by the the transmission rates, the evolution of the transmission mode is fundamentally governed by an eco-evolutionary feedback (see Figure 1).

**Figure 1:**
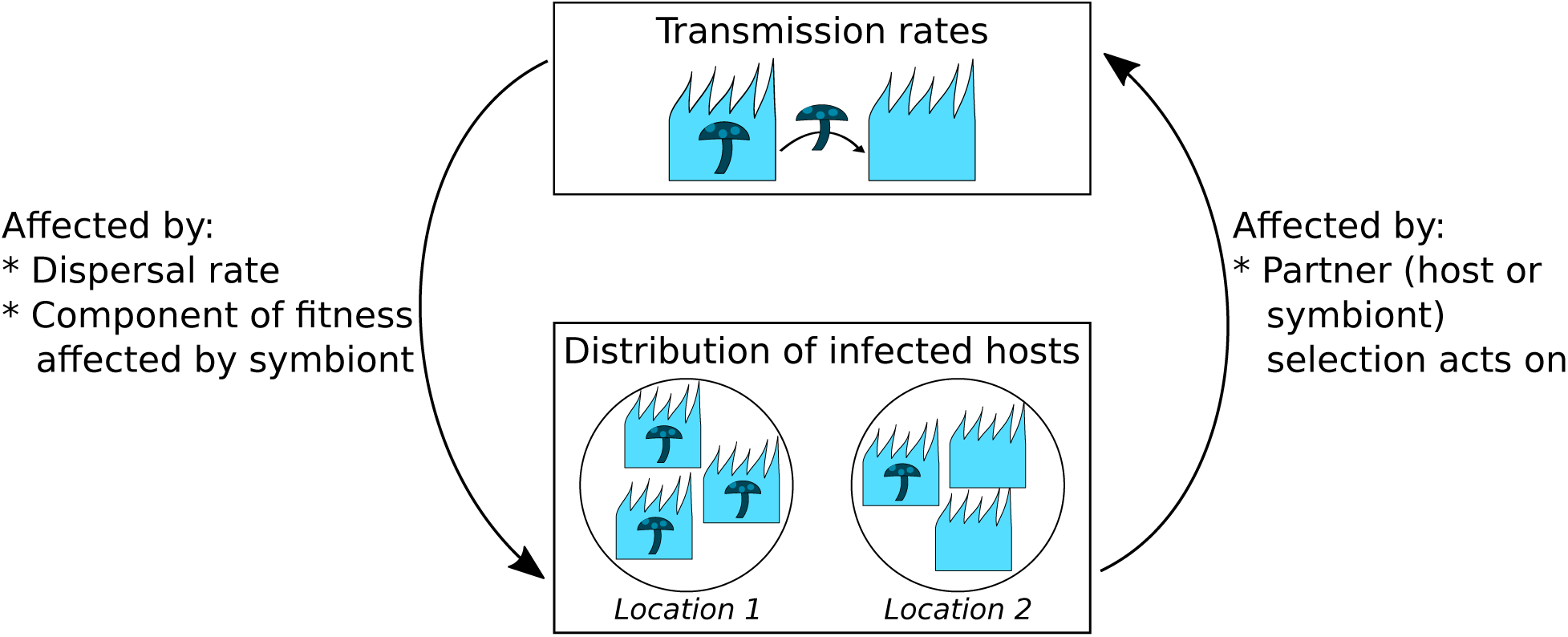
The evolution of transmission is governed by an eco-evolutionary feedback. The spatial distribution of infected hosts (bottom) affects the selective advantage of a mutant with a different transmission rate. As a mutant spreads, its transmission rates in turn influence the spatial distribution of infected hosts. The feedback from the spatial distribution to the transmission rates is influenced by the dispersal rate and the component of host fitness the symbiont affects. Similarly, the selective advantage of a mutant with new transmission rates is influenced by whether selection acts on the hosts or the symbiont, as different distributions of infected hosts are beneficial to them.

This eco-evolutionary feedback suggests that the evolution of transmission mode might ultimately depend on which life history stage is affected by the symbiont through the fitness component’s influence on the distribution of infected hosts. Accordingly, we find that when the symbiont affects host lifespan, a distribution of infected hosts that reflects the distribution of symbionts effects is produced at high horizontal transmission rates. This allows hosts evolve to high horizontal transmission rates. On the other hand, when the symbiont affects host fecundity, high horizontal transmission leads to high fractions of infected hosts in both patches. In this case, hosts generally evolve low horizontal transmission rates. Regardless of the type of symbiont effect, low host dispersal rates allow hosts with high vertical and low horizontal transmission rates to have high frequencies of infected hosts in locations where the symbiont is mutualistic and low frequencies elsewhere. This allows high vertical transmission to evolve at low but not high dispersal rates.

Our results highlight how ecological feedback from the fraction of infected hosts generated by the current transmission rates affects the selective advantage of mutant transmission rates, determining the course of evolution. This suggests that the manner in which the symbiont affects host life history and ecology ultimately influences host evolution and the ecological dynamics hosts evolve toward.

## 2 Methods

We first describe the model in general, then discuss the methods for the analytical and simulation models.

### 2.1 The Model

We model a patch-structured population where the symbiont is beneficial in half the patches (M-patches) and harmful in the other half (P-patches). We consider two types of conditional mutualism: one where the symbiont affects host fecundity and the other where it affects host lifespan. In the main text, we show results from the case where the symbiont affects host lifespan through the newborn host’s establishment probability. This is almost identical to the case where the symbiont affects lifespan through adult host mortality, which we show in the supplement, Figure S2. We analytically model the case where there are two patches of infinite size. For tractability in our analytical model, the ecological and evolutionary dynamics occur on separate timescales. We use simulations to investigate the effects of finite populations and concurrent ecological and evolutionary changes. In both cases, we assume all patches are of constant and equal size. We track the fraction of infected hosts in each patch (given by *i_q_* for patch *q*) and the horizontal and vertical transmission probabilities of the resident and mutant, (*h* and *υ* for the resident and *h*^*^ and *υ*^*^ for the mutant; see Table 1 for list of variables). We assume that neither multiple infection nor loss of the symbiont once infected is possible. When hosts control transmission, we assume that a host’s transmission probabilities determine its probability of infection. When symbionts control transmission, uninfected hosts cannot be said to have a transmission probabilities. Instead we model the potentially infecting symbiont as determining the transmission probability. Conflict over transmission mode might then occur between the host receiving the symbiont and the incoming symbiont.

**Table 1:**
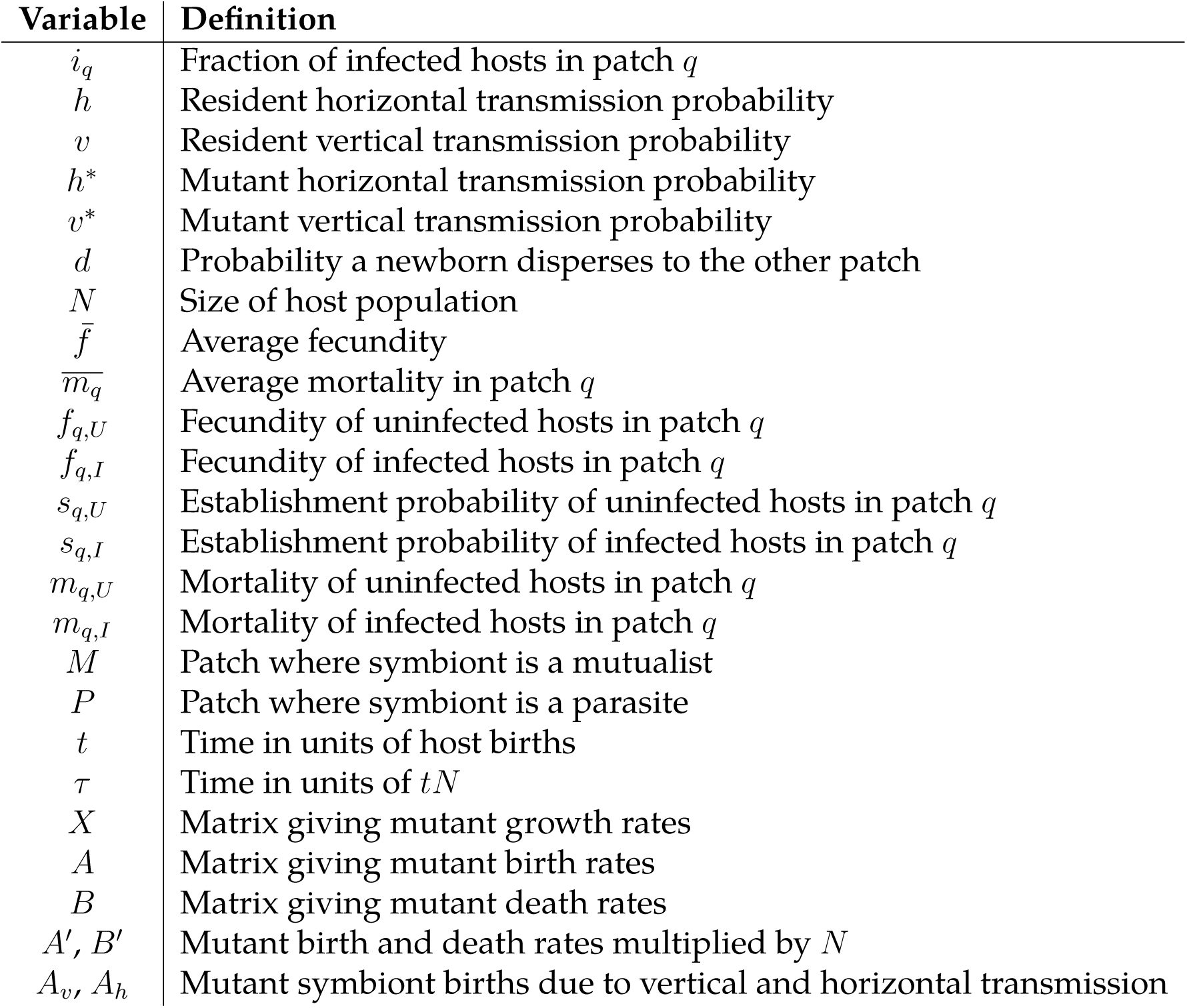
Variables

We model overlapping host generations in discrete time. The host lifecycle is given in Figure 2. Each time step a host is chosen to reproduce, with the probability of reproduction determined by the host’s patch and infection status. A host in patch *q* has fecundity *f_q,I_* if it is infected or *f_q,U_* if uninfected. The probability that a host with fecundity *f* reproduces is 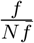, where *N* is the population size, and *f̅* is the average fecundity.

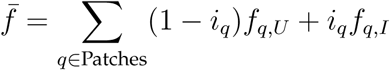

**Figure 2:**
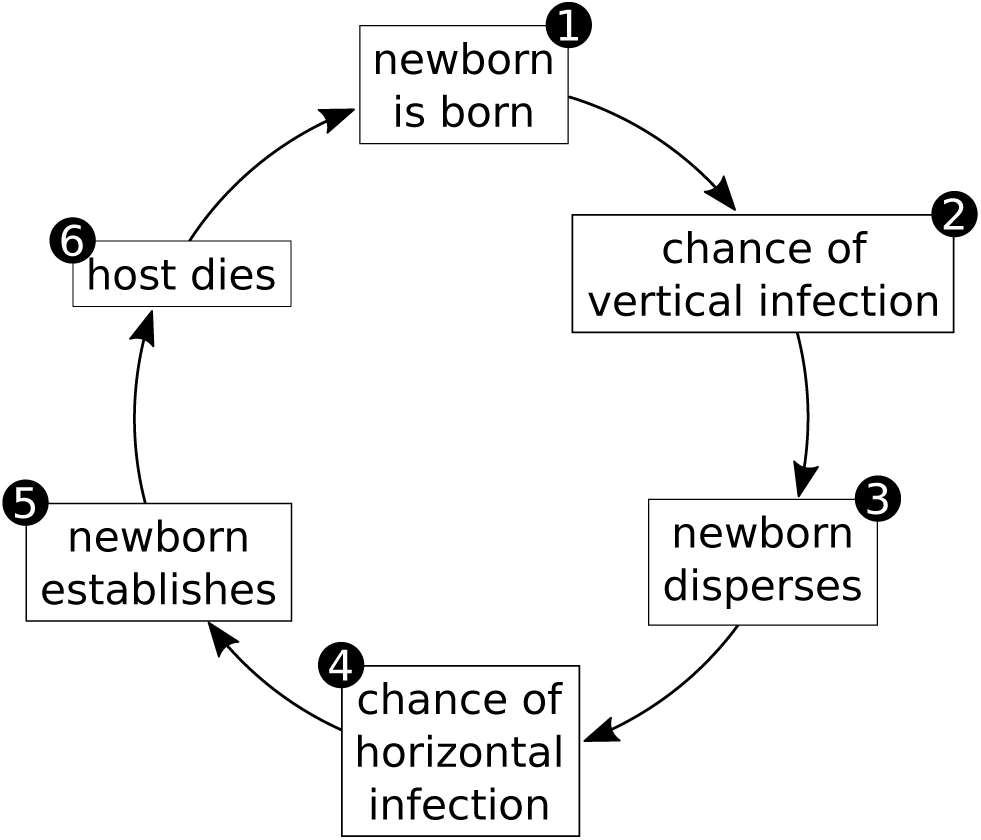
Host lifecycle. Numbers indicate the order the events happen in the simulations.

When the symbiont affects host fecundity, we assume infected hosts have higher fecundity than uninfected hosts in M-patches, and that the reverse is true in P-patches. When the symbiont affects host lifespan, we assume all hosts have equal fecundity.

If the parent host is infected, its offspring has a chance to acquire the symbiont via vertical transmission. For a vertical transmission probability *υ*, the probability that a host patch *q* gives birth to an uninfected or infected offspring is

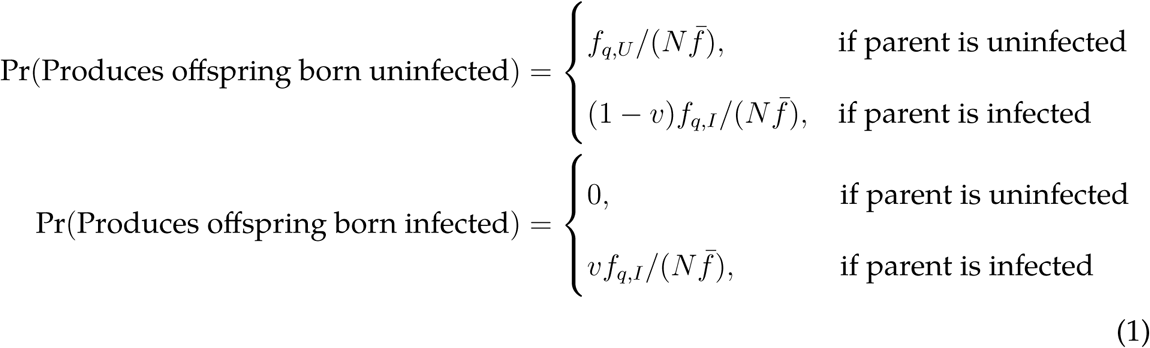

After birth, newborns disperse to a new patch with probability *d* or stay in their natal patch with probability 1 − *d*. We assume that newborns must mature somewhat before they become susceptible to horizontal infection, such that there is a window of time after dispersal and before establishment where newborns may acquire the symbiont horizontally, as is the case for many horizontally transmitted symbioses (Bright and Bulgheresi, 2010). For simplicity, we assume that when newborns arrive in the patch, they make contact with a single neighbor, who, if infected, may infect the newborn with probability *h*.

Once newborns have dispersed and become infected (or not), they must establish in their patch. Uninfected and infected newborns in patch *q* have establishment probabilities *s_q,U_* and *s_q,I_*, respectively. When the conditional mutualism affects host establishment, infected hosts are more likely to establish than uninfected in M-patches. The reverse is true in P-patches. When the symbiont affects fecundity, we set all establishment probabilities to 1 so that newborns always establish. (It would also be possible to assume all newborns have an establishment probability less than 1, but this makes the simulations slower without changing the results.)

For a newborn arriving in patch *q*, its chance of establishing as an uninfected (or in-fected) adult is

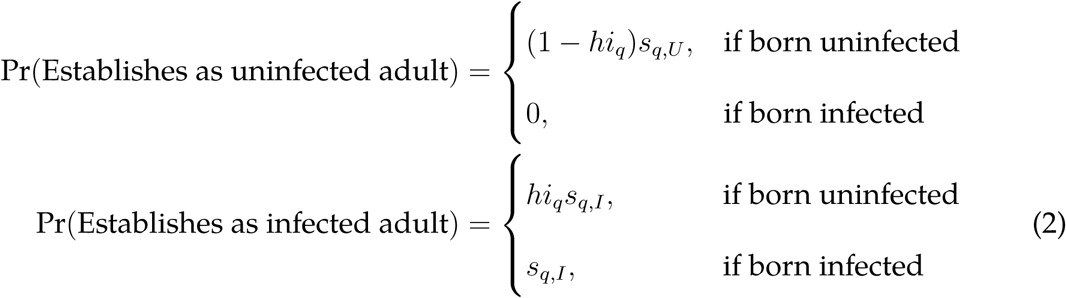

Finally, we assume patch sizes are constant, so if the newborn establishes, another host in the patch must die. If the newborn arrives in patch *q*, the probability that an adult host in *q* with mortality *m* dies is

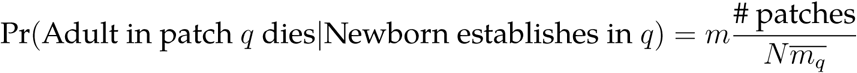

where *N* is the population size, and 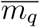 is the average mortality in patch *q*.

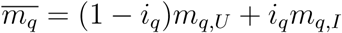

### 2.2 Analytical Model

Before we can determine the fitness of a mutant host or symbiont, we must know what fraction of hosts are currently infected in each patch. To determine the ecological equilibrium fraction of infected hosts in a monomorphic population, we find the point where the rate of change of the fraction of infected hosts in each patch vanishes. (The ecological equilibrium is not affected by whether hosts or symbionts control transmission evolution.) Assuming all fecundities and mortalities are nonzero, the rate of change of the fraction of infected hosts in patch *q* is

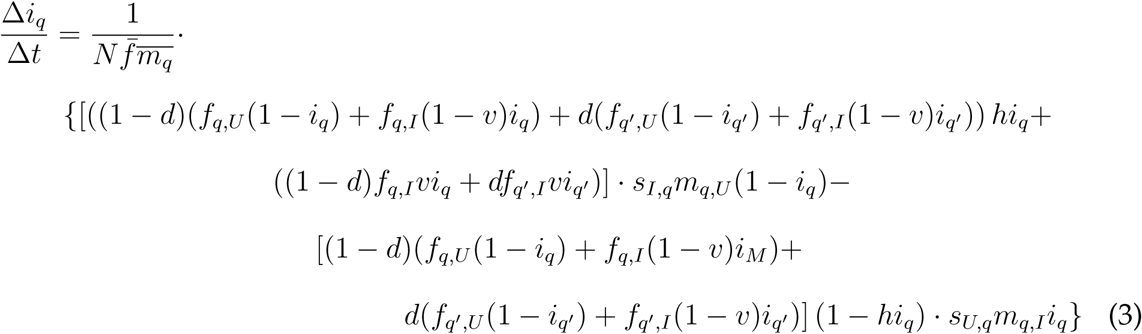

where *t* is the time in units of one host birth per time step (see Appendix A for derivation).

We use Mathematica version 11 (Wolfram Research Inc., 2017) to solve for the values of *i_M_* and *i_P_* that make Equation 3 be vanish for both patches (code given in supplement). While there may be multiple (*i_M_, i_P_*) pairs that satisfy the equation (for example (*i_M_* = 0*, i_P_* = 0) is always a solution), not all of them are stable in response to small perturbations in the fraction of infected hosts. We consider the monomorphic population ecological equilibria to be only those solutions that are stable in response to perturbations (see Appendix A). In most cases, there is only one stable ecological equilibrium. In cases where there is more than one ecological equilibrium, we show one equilibrium in the main text and the other in the supplement. In all cases that we investigated, multiple ecological equilibria for a given pair of transmission probabilities do not have qualitatively different effects on the overall pattern of transmission evolution.

To determine the direction transmission rates evolve in, we find the invasion fitness of a mutant with slightly different horizontal and vertical transmission probabilities than the resident. Because mutants in different patches (and, for mutant hosts, mutants with different infection statuses) differ in their chances of producing offspring, we model the growth of the mutant when rare as a multitype branching process (Lehmann et al., 2016). We write a matrix *X_τ_* that gives the expected number of mutants produced by a mutant in each patch (or, for host control, an uninfected or infected mutant in each patch) at every time step, measuring time in units of host births times population size, *τ* = *tN*. The leading eigenvalue of *X_τ_* then gives the growth rate of the mutant when rare. The derivation of *X_τ_* for host and symbiont control follows straightforwardly from Equations 1 and 2 and is given in detail in Appendix A.

Once we have *X_τ_*, we can calculate the derivative of the mutant growth rate in terms of the mutant transmission probabilities. We can then use these derivatives to trace the path of transmission evolution. We find the derivatives of the leading eigenvalue of *X_τ_* numerically and then numerically calculate the path of the evolutionary trajectories in Mathematica (see Appendix A).

### 2.3 Simulations

We simulate transmission mode evolution in Julia version 0.5.1 (Bezanson et al., 2017, the simulation code is available as a supplementary material). Each time step, events happen in the order in Figure 2, starting from host birth. A single host is selected to give birth, with the probability of selection determined by its patch and infection status. After a host is born, if hosts control transmission, we allow the newborn’s transmission probabilities to mutate. In the case of host control, the newborn host’s possibly mutated new transmission probability determines its probability of infection. When symbionts control transmission, the parent’s symbiont determines the vertical transmission probability, and then if infection is successful, the newborn’s symbiont is allowed to mutate.

The newborn then disperses to a new patch with probability *d* and remains in its natal patch with probability 1 − *d*. If the newborn disperses, it is equally likely to end up in any patch except its natal one. If the newborn is so far uninfected, a random adult host in the newborn’s patch is then selected to be its potentially infection contact. If this adult is infected, horizontal transmission occurs with probability given by the newborn’s horizontal transmission probability (host control case) or the neighbor’s symbiont’s horizontal transmission probability (symbiont control case). If the newborn becomes infected and the symbiont controls transmission, the newborn’s symbiont may then mutate. Finally, the newborn’s establishment in the patch is determined by its infection status and location. If the newborn successfully establishes, a random adult host is chosen to die.

Before allowing transmission mode to evolve, we ran the simulation for 4000 time steps to allow the resident population to equilibrate. We started the simulations from an 11×11 grid starting points evenly spaced over the space of all possible transmission probabilities. After the equilibration period, we ran each simulation for 10^7^ time steps. We used a mutation rate of 0.02, with mutations normally distributed with a mean of the originally transmission probability and standard deviation of 0.05. For the host control case, we also had a 0.5% chance that an uninfected newborn would be spontaneously infected. We did this to prevent the infection from being lost by chance leading transmission to evolve neutrally for the rest of the simulation. We analyzed the simulations by finding the average transmission rates and fraction of infected hosts in M- and P-patches at the last time step using the plyr package (Wickham, 2011) in R (R Core Team, 2017).

## 3 Results

### 3.1 Host Control of Transmission

The results for infinite populations suggest that different factors control when vertical and horizontal transmission can evolve. Vertical transmission evolves when newborn hosts rarely disperse from their natal patch. Horizontal transmission evolves when there is a higher fraction of infected hosts in M-patches than P-patches. When the symbiont affects fecundity, high horizontal transmission erodes the difference in the fraction of infected hosts between patches. The difference is maintained when the symbiont affects lifespan. High horizontal transmission is more likely to evolve when the symbiont affects lifespan. When we simulate finite populations, polymorphism in transmission probabilities between patches arises at low dispersal rates. At high dispersal rates, the simulations resemble the infinite population case.

#### 3.1.1 Analytical Model (Infinite Population)

##### Symbiont Affects Lifespan

When the symbiont affects host lifespan in a monomorphic population, the ecological equilibrium fraction of infected hosts is generally higher in M-patches than P-patches (Figure 3a-f), except when both transmission probabilities are too low and the infection dies out (white regions in Figure 3) or when both transmission probabilities are 1 and all hosts in both patches are infected. In both cases, transmission evolves neutrally, since changes in transmission do not affect a host’s chances of becoming infected.

**Figure 3:**
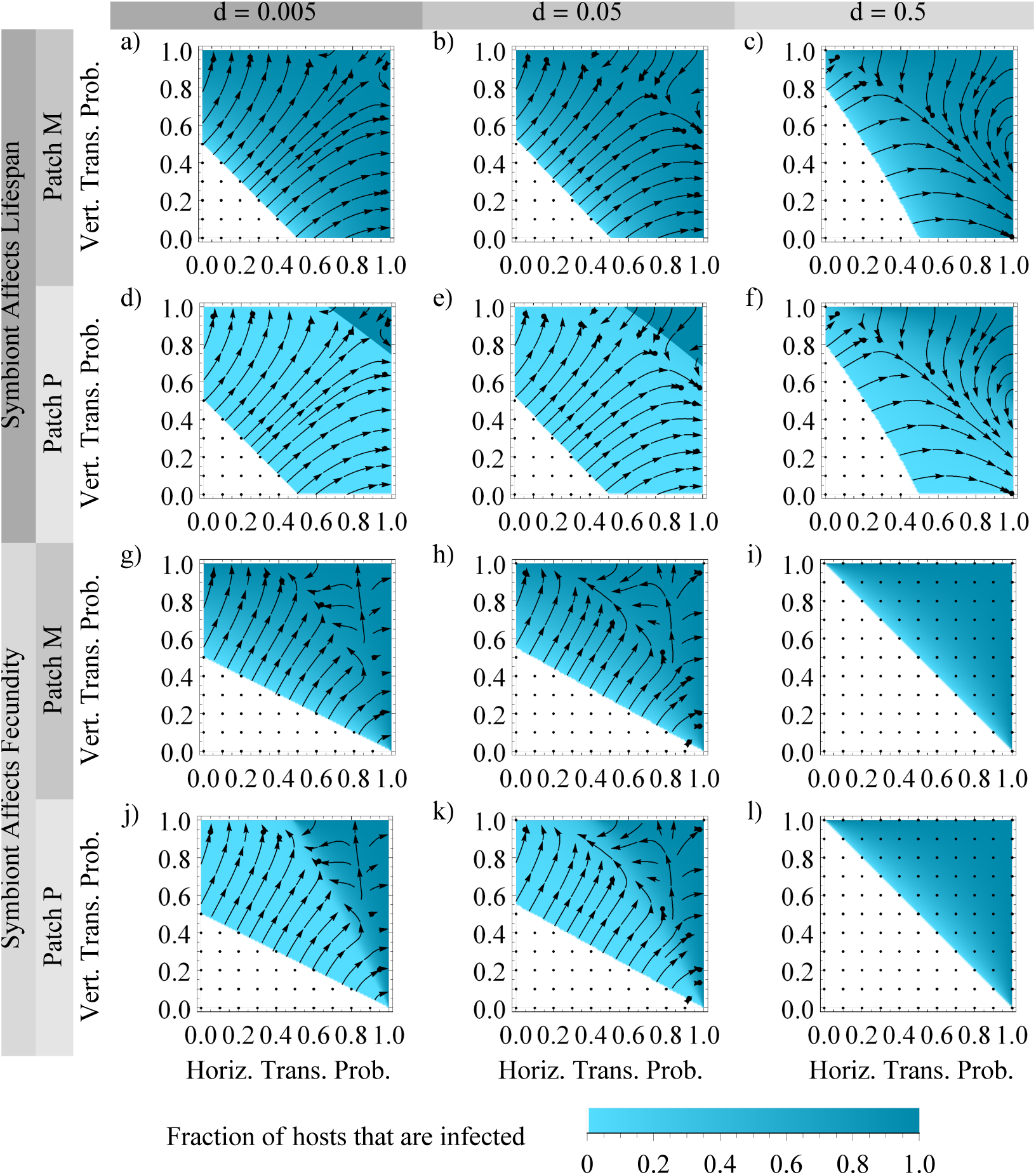
Ecological equilibria and host evolutionary trajectories for an infinite population. Panels a-f: symbiont affects host lifespan, panels g-l: symbiont affects host fecundity. Columns indicate dispersal rates. The upper and lower pairs of panels in a column each represent a single metapopulation, with the upper panel indicating the fraction of infected hosts in Patch M, and the lower the fraction of infected hosts in Patch P (e.g. panels a and d represent a single population). For each plot, colors indicate the fraction of infected hosts in the patch when the population is monomorphic for a given pair of horizontal and vertical transmission rates. Arrows indicate hosts evolutionary trajectories, with dots where transmission evolves neutrally. Panels from the same metapopulation show the same trajectories, as the entire population evolves together. Parameters, panels a-f: *f_M,U_* = *f_P,I_* = 0.5, *f_M,I_* = *f_P,U_* = 1, *s_M,U_* = *s_M,I_* = *s_P,U_* = *s_P,I_* = 1; panels g-l: *f_M,U_* = *f_P,I_* = *f_M,I_* = *f_P,U_* = 1, *s_M,U_* = *s_P,I_* = 0.5, *s_M,I_* = *s_P,U_* = 0.5.

Aside from the above cases, host evolutionary trajectories lead to either complete horizontal and no vertical transmission, i.e. (*h* = 1, *υ* = 0); or they lead to complete vertical transmission and no horizontal transmission, (*h* = 0, *υ* = 1). At low dispersal rates, the basins of attraction of the two endpoints are very similar in size (Figure 3a,d). As the dispersal rate increases, more trajectories lead to the point (*h* = 1, *υ* = 0). This corresponds to changes in the transmission probabilities that lead to high fractions of parasitized hosts. As dispersal increases, even intermediate values of horizontal and vertical transmission paired with high levels of the other lead to a large fraction of infected hosts in Patch P. However, the effect is more pronounced for high vertical transmission probabilities, which require much lower horizontal transmission probabilities in order to contain the symbiont to Patch M. (This can be seen in the increasing length of the top of the dark triangle in Figure 3d-f compared to its right side.) Finally, when the dispersal rate is maximum (*d* = 0.5 for the two patch case, meaning newborns have an equal chance of ending up in either patch), all host evolutionary trajectories lead to complete horizontal and no vertical transmission (Figure 3c,f). This is because high vertical transmission leads to a high fraction of parasitized hosts for all horizontal transmission probabilities, including *h* = 0.

While the basin of attraction of high horizontal versus high vertical transmission depends on the dispersal rate, evolutionary trajectories always lead to a beneficial (to hosts) distribution of the symbiont, in the sense that they maintain a high fraction of infected hosts in the patch where the symbiont is mutualistic and a low fraction of infected hosts in the patch where the symbiont is parasitic.

##### Symbiont affects fecundity

When the symbiont affects fecundity, high horizontal transmission probabilities always lead to a high ecological equilibrium fraction of infected hosts in Patch P. In contrast, high vertical transmission probabilities, combined with low horizontal transmission probabilities, produce the largest difference in the fraction of infected hosts between Patches M and P (Figure 3g-l). As a result, most trajectories lead to complete vertical and no horizontal transmission, (*h* = 0, *υ* = 1).

However, unlike the case where the symbiont affects lifespan, not all trajectories lead to transmission probabilities that contain the symbiont to the patch where it is beneficial. When dispersal is not maximum (*d* < 0.5), populations that start with too high horizontal transmission probabilities evolve towards complete infection, due the fact that symbionts become abundant everywhere, and therefore the host has little chance of escaping them in Patch P by a small decrease in transmission rates. Therefore, there is little additional cost to hosts from increasing transmission in Patch P, and a slight benefit in Patch M. Trajectories that lead to complete infection end up in one of two regions. In the first region, the population has complete horizontal transmission and at least some vertical transmission, (*h* = 1, *υ* > 0). In the second region, the population has complete vertical transmission and high horizontal transmission, (*h* > *h*^*^ *υ* = 1). The precise value of *h*^*^ depends on the dispersal rate and the costs/benefits provided by the symbiont. Interestingly, if the symbiont is more costly in Patch P than it is beneficial in Patch M, all trajectories lead to the point (*h* = 0, *υ* = 1). On the other hand, if the symbiont is more beneficial in Patch M than harmful in Patch P, populations are more likely to evolve towards complete infection (Figure S3).

As the dispersal rate increases, a relatively high frequency of parasitized hosts appear at increasingly lower values of horizontal transmission, particularly at high vertical transmission probabilities (as shown by the increasing size of the dark regions at the top of Figure 3 from panels j to k). This means that more evolutionary trajectories start in regions where the symbiont is not well contained to Patch M, and a small decrease in transmission probabilities is not as beneficial to hosts in Patch P as an increase is to hosts in Patch M. More trajectories therefore lead to complete infection in both patches.

Finally, when newborns have an equal chance of ending up in either patch (dispersal rate = 0.5, Figure 3i-j), the two patches have the same frequency of infected hosts at all transmission probabilities. When the symbiont’s costs in Patch P exactly equal its benefits in Patch M (as in Figure 3), transmission is selectively neutral. The benefits of a small increase or decrease in one patch are exactly balanced with the cost of that change in the other. If the costs and benefits are not equal (Figure S3), hosts will either evolve towards low transmission and loss of the symbiont (when the costs are higher than the benefits) or high transmission and complete infection (when the benefits are higher than the costs).

### 3.2 Symbiont Affects Lifespan and Fecundity

In the supplement, we investigate the case where the symbiont affects both host lifespan and fecundity. In general, if the symbiont’s effect on one fitness component is significantly stronger than the other, transmission evolution largely resembles the case where only the stronger effect is present (Figures S4 and S5). One exception is if the symbiont largely affects fecundity and the dispersal rate is maximum. When the symbiont affects fecundity equally in both patches and does not affect lifespan, transmission mode is selectively neutral when dispersal is maximum. However, a small symbiont effect on lifespan can break the symmetry and allow hosts to evolve toward either complete infection, loss of the symbiont, or even the point (*h* = 1, *υ* = 0). (The last of these provides a small degree of symbiont containment.)

When the symbiont has a strong effect on both components of host fitness, the results are more complicated. The outcome depends on the conditions which trigger the effects on each component as well as the relative strengths of the effects on each component. However, two general trends emerge. The first is that using high horizontal combined with low vertical transmission to contain the symbiont to M-patches is only an option when the symbiont can decrease lifespan. For example, when the symbiont affects fecundity, adding a conditional (in Patch P) or unconditional (in Patches M and P) lifespan cost to infection allows horizontal transmission to evolve as a method of containment (Figures S4 and S6).

Related to this, symbiont containment can often be improved by increasing the costs of infection. If trajectories do not lead to containment, increasing the cost of infection through fecundity or lifespan effects, can increase the number of trajectories leading to symbiont containment (Figures S4, S6, and S7). This is true even if hosts in M-patches bear the additional cost of infection (Figures S6 and S7). (On the other hand, increasing the cost of infection can also cause the symbiont to be lost in some cases, generally when the dispersal rate is maximum and the symbiont largely affects fecundity, e.g. Figure S6.)

#### 3.2.1 Simulations (Finite Population)

At high dispersal rates, the simulations of finite host populations behave much like the infinite population case (Figure 4c,f,i,l). However, as the dispersal rate decreases, the simulations diverge from the analytical results, in that the patches behave more like separate populations. At low dispersal rates, hosts residing in the patch where the symbiont is beneficial have higher average transmission probabilities than predicted for the infinite population case (Figure 4a,g have a large proportion of simulations with high average horizontal and vertical transmission probabilities, while the infinite population case predicts only one high transmission probability). Patches where the symbiont is parasitic tend to lose the infection (or have the symbiont at very low frequencies due to spontaneous infection) and then have transmission probabilities that evolve neutrally (Figure 4d,j and Figure S8). As the population size increases, lower dispersal rates are needed for the population to behave like separate patches, and the population resembles the infinite population at increasingly lower dispersal rates (Figure S9).

**Figure 4:**
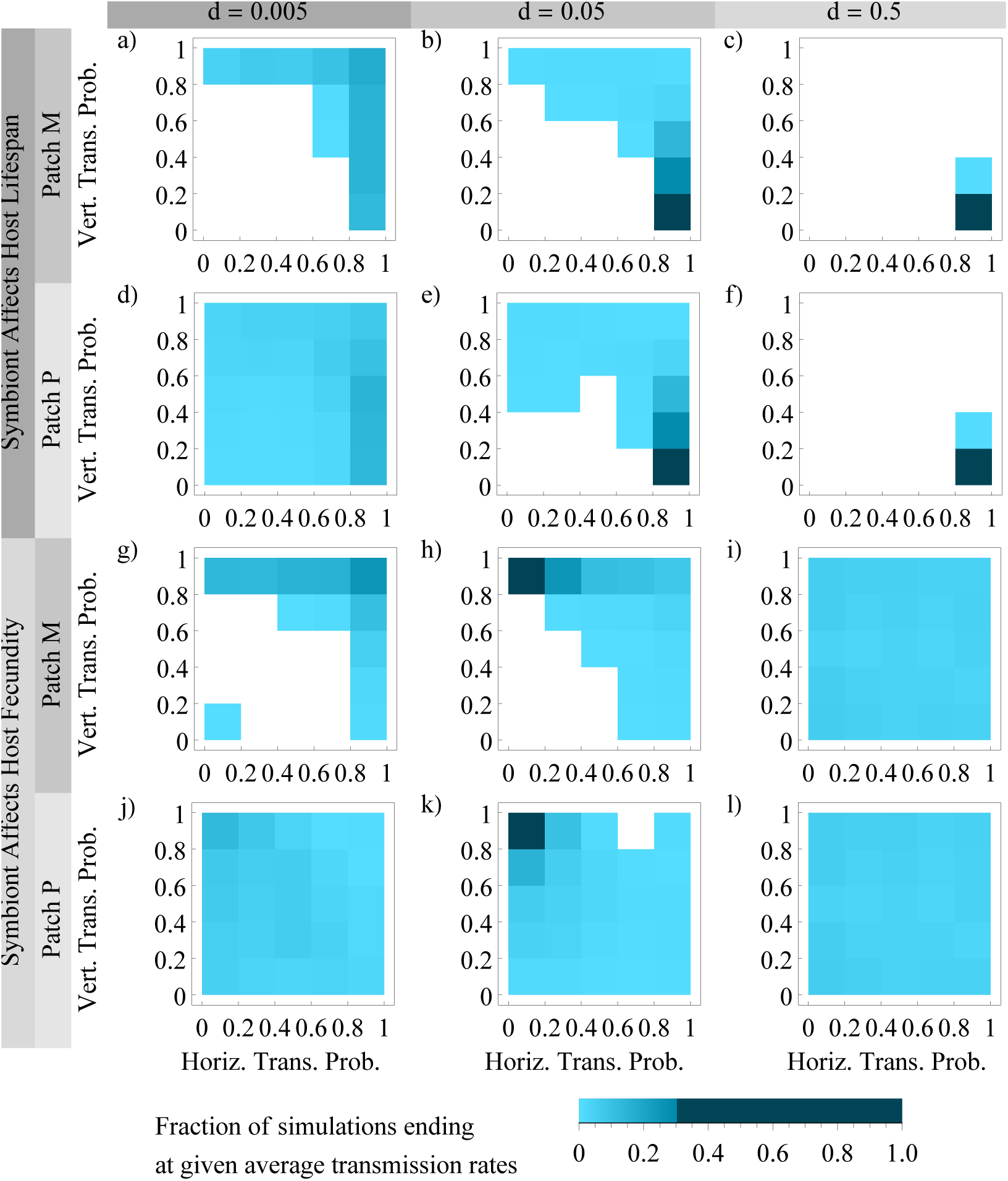
Simulations of transmission evolution under host control. Colors indicate fraction of populations ending with each combination of average horizontal and vertical transmission probabilities. Simulations were started from a grid of start points spaced 0.1 apart in transmission probability. Ten simulations at each start point were run for 10^7^ time steps for every parameter combination. Parameters: 2 patches, *N* = 200, mutation rate = 0.02, mutation standard deviation = 0.05, spontaneous infection probability = 0.005, panels a-f: *f_M,U_* = *f_P,I_* = 0.5, *f_M,I_* = *f_P,U_* = *s_M,U_* = *s_M,I_* = *s_P,U_* = *s_P,I_* = 1, panels g-l: *s_M,U_* = *s_P,I_* = 0.5, *f_M,U_* = *f_M,I_* = *f_P,U_* = *f_P,I_* = *s_M,I_* = *s_P,U_* = 1.

### 3.3 Symbiont Control

In both the analytical model and simulations, symbionts evolve high horizontal and vertical transmission probabilities (Figures S10 and S11). In particular, symbionts always evolve complete vertical transmission in the infinite population case. The horizontal transmission probability evolves neutrally once 100% vertical transmission is reached. This may be due to the fact that at high levels of infection, vertical transmission guarantees access to uninfected hosts to infect. The difference between selection pressure on hosts and symbionts is shown in Figure 5. In general, the most conflict is found at high vertical transmission probabilities. When the symbiont affects lifespan, conflict occurs at high vertical and horizontal transmission. As the dispersal rate increases and vertical transmission becomes less beneficial to hosts, the region of conflict expands to include low vertical transmission and intermediate transmission. This creates a triangular region where too much transmission, and particularly too much vertical transmission, leads to host-symbiont conflict. When the symbiont affects host fecundity, most conflict still occurs at high vertical transmission probabilities, but now intermediate horizontal transmission provokes the most conflict. This is because hosts at high horizontal transmission probabilities evolve towards complete infection, reducing the conflict between hosts and symbionts.

**Figure 5:**
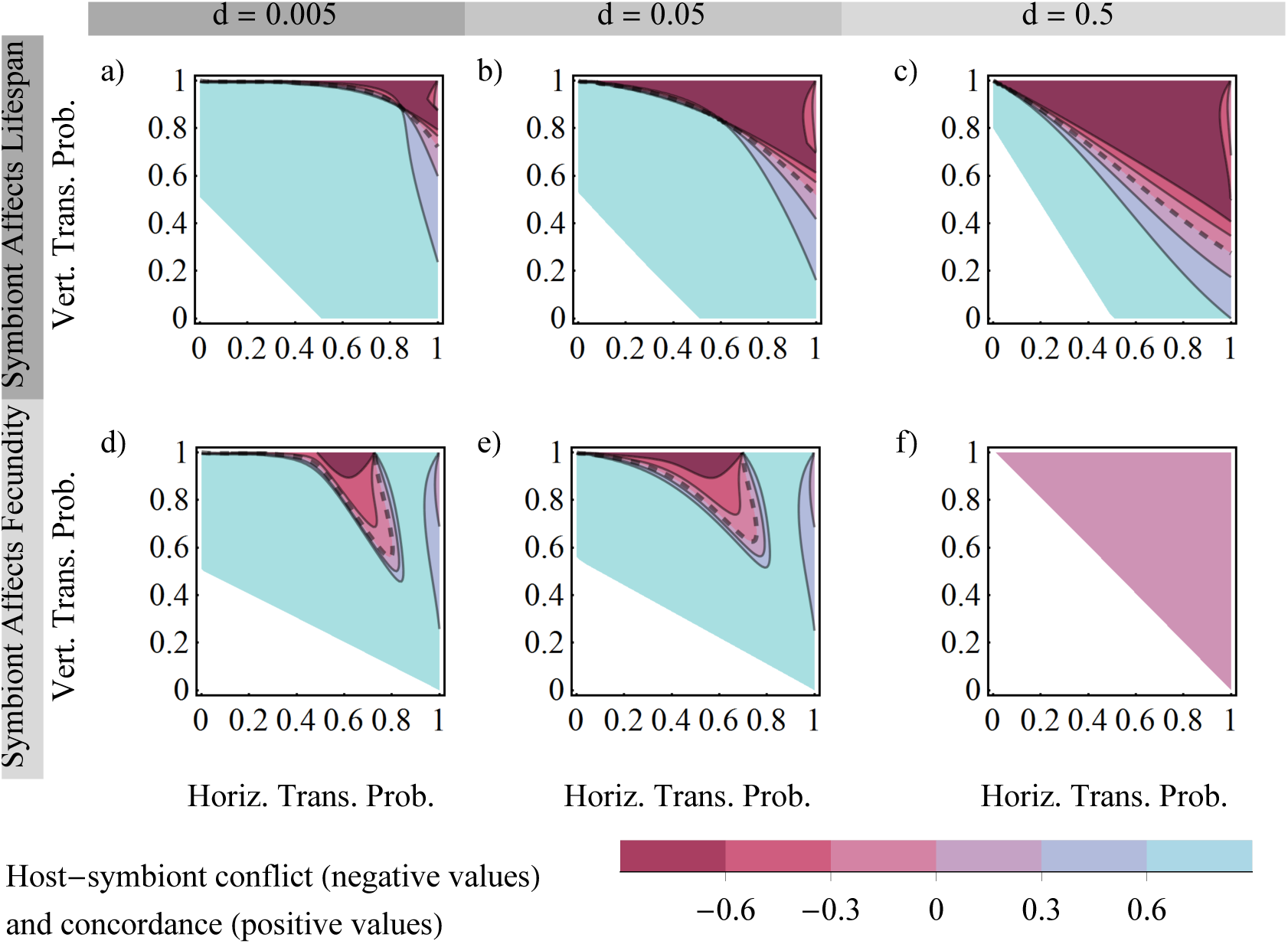
*Host-symbiont conflict:* Host-symbiont conflict when symbiont affects lifespan (top row) or fecundity (bottom row). Colors indicate the degree to which host and symbiont evolutionary trajectories point in the same direction, defined as the defined as the cosine of the selection vectors under host and symbiont control, or 0, if at least one of the selection vectors has magnitude 0. If trajectories are perpendicular or a partner does not experience selection, conflict is 0. Negative values indicate trajectories point in opposite directions (conflict), and positive values indicate that trajectories point in the same direction (concordance). Dashed lines separates regions of conflict and concordance. White regions indicate transmission rates where the infection cannot be maintained. Parameters, top row: *f_M,U_* = *f_P,I_* = 0.5, *f_M,I_* = *f_P,U_* = 1, *s_M,U_* = *s_M,I_* = *s_P,U_* = *s_P,I_* = 1; parameters, bottom row: *f_M,U_* = *f_M,I_* = *f_P,U_* = *f_P,I_* = 1, *s_M,U_* = *s_P,I_* = 0.5, *s_M,I_* = *s_P,U_* = 1.

## 4 Discussion

We investigate conditional mutualisms with spatial variation in symbiont quality and find that hosts evolve different transmission modes depending on the ecological distribution of infected hosts, which in turn depends on the aspect of fitness symbionts affect. When symbionts affect host lifespan, hosts are able to evolve high horizontal and low vertical transmission, which contains the symbiont to the patch where it is a mutualist. They are able to do this because hosts with the “wrong” status die more quickly and do not remain in the population to affect incoming newborns’ chance of infection. This sets up a difference in the distribution of infected hosts so that newborns benefit from higher horizontal transmission rates, because their probability of acquiring the symbiont is higher where it is beneficial.

When the symbiont affects fecundity, hosts with the “wrong” infection status reproduce less, but remain in the population just as long, which allows them to affect the infection status of incoming newborns. Unless the distribution of infected hosts is already skewed toward more infected hosts in the patch where the symbiont is beneficial, hosts gain no benefit from evolving horizontal transmission. Even worse, an increase in horizontal transmission produces some hosts with the “wrong” infection status, who then persist in the population to alter the infection probabilities of incoming newborns. This means that past a threshold transmission probability, horizontal transmission is no longer effective at maintaining different distributions of infected hosts. Hosts are left with using vertical transmission to contain the symbiont when dispersal is low and host lineages are mostly confined to the same patch. When dispersal is at its maximum, the patches have equal fractions of infected hosts, and the costs and benefits of infection determine if the infection is lost (when the symbiont is more harmful in P-patches than beneficial in M-patches), spreads to everyone (when the symbiont is less harmful in P-patches than beneficial in M-patches), or drifts because transmission rate is neutral (when symbiont costs and benefits are exactly equal).

When the symbiont affects lifespan and fecundity, the nature and magnitude of the costs of infection have a large influence on transmission evolution. Hosts are only able to use horizontal transmission to contain the symbiont when the symbiont decreases lifespan. This decrease in lifespan does not have to be conditional on hosts’ environment in order to allow symbiont containment. Furthermore, adding conditional or unconditional lifespan or fecundity costs of infection can increase the fraction of host evolutionary trajectories that lead to symbiont containment, rather than complete infection. These results suggest that the costs of a conditional mutualism are key to determining its evolutionary outcome. They also suggest that a conditional mutualism that has more costs than benefits may actually be better for hosts than more “mutualistic” conditional mutualisms, by increasing hosts’ chances of evolving transmission modes that contain the symbiont to locations where it is beneficial.

The simulations largely confirm that our results hold for finite populations. However, they suggest an alternative way that hosts in small populations may respond to a conditional mutualism when dispersal rate is low. If dispersal rate is small enough relative to the population size, the subpopulations of hosts in each patch behave more like separate populations, and exhibit local adaptation. Hosts in M-patches evolve high horizontal and vertical transmission rates, while hosts in P-patches lose the symbiont (or have it at low frequency due to spontaneous infection) and have transmission evolve neutrally. This suggests that at low dispersal rates, it is possible that hosts in small populations have more options for transmission mode evolution. Hosts whose symbiont affects their fecundity may not be constrained to use purely vertical transmission when the dispersal rate is low. However, the main problem for hosts still occurs at high dispersal rates, when the patches do not behave like separate populations, and hosts whose symbiont affects fecundity are forced to have the same fraction of infected hosts in both patches. As it is unlikely in nature that symbiont costs and benefits will be exactly balanced, in practice this may lead to the symbiont being lost if it is slightly more harmful or maintained in all hosts if it is slightly more beneficial.

Our model of symbiont control shows that, as predicted, when there are no direct costs to transmission and population size is fixed, symbionts evolve high transmission rates and end up infecting all hosts in the population. In both the analytical and simulation models, symbionts evolve complete vertical transmission and evolve a nonzero probability of horizontal transmission that guarantees complete infection of all hosts (this may be less than a 100% chance of horizontal transmission, since vertical transmission also contributes to the chance of infection). Further, vertical, rather than horizontal, transmission is maximized because at high frequencies of infected hosts, vertical transmission is the best way to guarantee that newborns are infected (Lipsitch et al., 1995).

Our results can be used to predict the spread of symbionts and transmission mode evolution in known conditional mutualisms, if the symbiont’s effect on the host and the dispersal rate are known. For example, in the symbiosis between aphids and their obligate symbiont *Buchnera aphidicola*, a mutation in the promoter of *ibpA*, which encodes one *B. aphidicola*’s heat shock proteins, causes mutant *B. aphidicola* to increase host fecundity (relative to wild-type *B. aphidicola*) in cool conditions and nearly eliminate reproduction in warm conditions (Dunbar et al., 2007). The mutant has been found at frequencies up to 20% in natural populations, despite its large potential cost and the fact that *B. aphidicola* is strictly vertically transmitted. Our results suggest that the lack of horizontal transmission is not necessarily a barrier to the persistence of the symbiont in natural populations, and may in fact benefit its hosts, provided that aphid dispersal between regions with different temperatures in relatively rare.

One other example to which we can apply our model is the symbiosis between the grass *Agrostis hyemalis* and the fungus *Eplichloë amarillans*. *E. amarillans* increases host fecundity under drought conditions and decreased host biomass in the presence of certain soil microbes (Davitt et al., 2011). It is difficult to know exactly how biomass affects lifespan and fecundity, but as long as biomass has a smaller effect on lifespan than fecundity, we would predict that vertical transmission, particularly if seeds disperse to new environments only rarely, would be more likely to arise. Indeed, vertical transmission is observed in this symbiosis, although without knowing the relative effect of biomass on lifespan and fecundity, it is difficult to be certain whether the system matches our predictions.

While many other conditional mutualisms are known, in most of these the symbiont’s effect on different components of host fitness is currently unknown. Our results suggest that quantifying context-dependent variation in fitness components could allow predictions of transmission mode evolution and symbiont spread.

An important overall conclusion from our model is that in conditional mutualisms, it is not just the costs and benefits of infection that matter, but also the component of fitness that the symbiont affects. The component of fitness influences the distribution of the infection on ecological timescales, meaning it may be useful for predicting the spread of conditional mutualisms of interest. The ecological distribution of infected hosts also strongly influences transmission mode evolution. As transmission mode is predicted to itself create selective pressure on virulence, the ecological distribution of infected hosts over evolutionary time may feed back not only on transmission but also on the nature of the symbiosis itself. Thus, the feedback we found between symbiont effects on host fitness and transmission evolution may be important for predicting both the short- and long-term future of conditional mutualisms. As more symbiosis are being found to have conditional effects, understanding the precise nature of symbiont effects on their hosts may be useful for predicting the short-and long-term future of these symbioses.

## Appendix

### A Calculations for Infinite Population Model

#### A.1 Equilibrium Distribution of Infected Hosts

From Equations 1 and 2, we can see that the fraction of infected hosts in a patch affects hosts’ birth, establishment, and death probabilities, as well as symbionts’ transmission opportunities. So, before we can find the invasion fitness of a mutant host or symbiont, we need to find the equilibrium fraction of infected hosts. We find the equilibrium fraction of infected hosts analytically for an infinite host population with two patches. We call these patches M and P and assume they are each of size 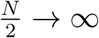. In patch M, the symbiont is a mutualist that increases either infected host fecundity or lifespan (depending on the nature of the conditional mutualism) above that of uninfected hosts. In patch P, the reverse is true. We will usually assume either *f_M,I_* = *f_P,U_* > *f_M,U_* = *f_P,I_* or *s_M,I_* = *s_P,U_* > *s_M,U_* = *s_P,I_*. In the supplement, we relax this assumption and also consider the case where the symbiont affects lifespan through adult mortality (*m_M,I_* = *m_P,U_* > *m_M,U_* = *m_P,I_*).

To find the equilibrium fraction of infected hosts in patches M and P, we must solve

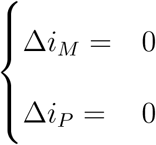

for the fraction of infected hosts in each patch, *i_M_* and *i_P_*.

To do this, we must write down formulas for the change in infected hosts in a patch. The fraction of infected hosts in a patch should increase if an infected newborn establishes and an uninfected adult dies. It should decrease if an uninfected newborn establishes and an infected adult dies. All other events (newborn failing to establish, uninfected newborn establishing in place of an uninfected adult, infected newborn establishing in place of an infected adult) should not lead to a change in the frequency of infected hosts in the patch.

Because each patch is of size 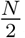, the addition or subtraction of a single infected host should change the frequency of infected hosts in the patch by 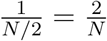. The rate of change in frequency in infected hosts in a patch should then be

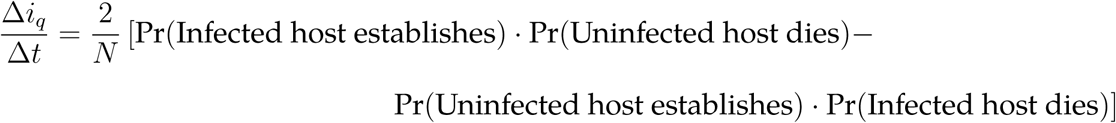

where *t* is time in units of host births, such that one host is born every time *t* increases by 1.

Using Equations 1 and 2, and taking into account the fact that newborn hosts may enter a patch via dispersal, the rate of change in the fraction of infected hosts is

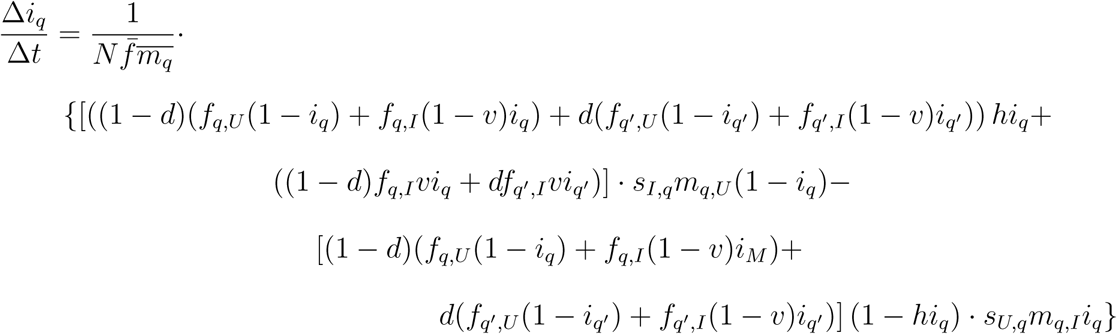

where *q* represents patch *M* or *P*, and *q′* is the other patch. Note that the rate of change is now scaled by 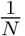, because there are 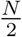 hosts in the patch which each have their chance to reproduce scaled by 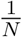.

By constraining all fecundities and mortalities (*f_M,U_*, *m_M,U_* etc.) to be greater than 0, we can ensure that the average fecundity, *f̄* and both average mortalities, 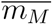 and 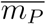 are always greater than 0. Then we can solve the slightly simpler set of equations

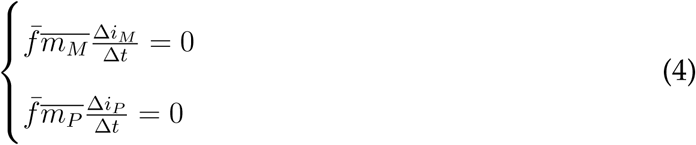

We solve this system numerically in the supplement using Mathematica version 11.1 (Wolfram Research Inc., 2017).

It is possible that some of the equilibrium fractions of infected hosts may not be stable. To find stable equilibria, we select those solutions of equation 4 for which the eigenvalues of the Jacobian are negative. The Jacobian is defined as

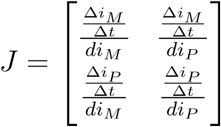

We find the eigenvalues of the Jacobian at each equilibrium numerically using Mathematica (supplement) and select those equilibria that are stable for invasion analysis.

#### A.2 Transmission Mode Evolution - Host Control

We can now investigate transmission mode evolution when transmission is a host trait. We want to find the invasion fitness of a mutant host with slightly different horizontal and vertical transmission rates than the resident. To do this, we can think of the growth of the mutant when rare as a multitype branching process (Lehmann et al., 2016). We write a matrix (*X_t_*) that gives the expected number of mutants produced by an uninfected or infected mutant in each patch every time step (measuring time in units of host births, *t*). Rows of *X_t_* correspond to the location and infection status of mutants produced. The first two rows correspond to uninfected and infected mutants produced in patch M, and the third and fourth rows are the same for patch P. Columns of *X_t_* correspond to the type of mutant producing a new mutant (or “producing” itself by surviving to the next time step). Columns are in the same order as rows. Then we have

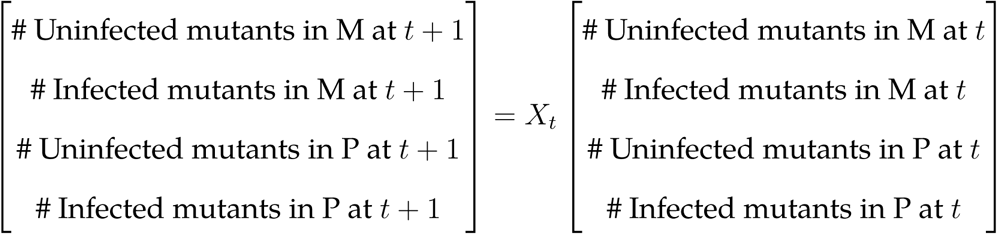

To find *X_t_*, let *A* be a matrix that gives the probability a mutant gives birth to an uninfected or infected offspring that successfully establish in each patch (rows and columns in same order as in *X_t_*). Let *B* be a matrix that gives the probability that an uninfected or infected mutant in each patch dies. Then

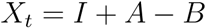

where *I* is the identity matrix and indicates that besides giving birth and dying, mutants may simply persist in the population from one time step to the next.

We can get the probabilities in *A* from the product of Equations 1 and 2. The probabilities we need for *A* are the following:

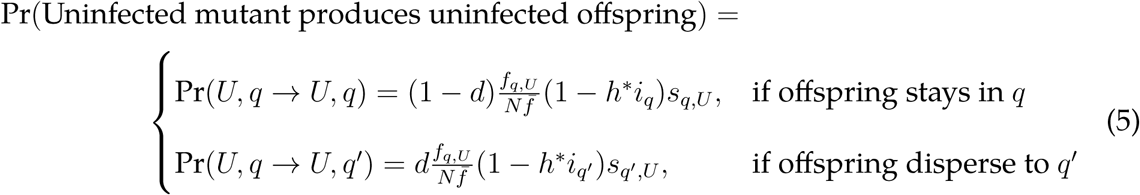

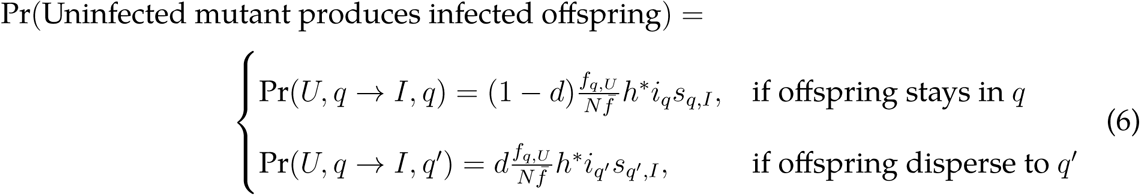

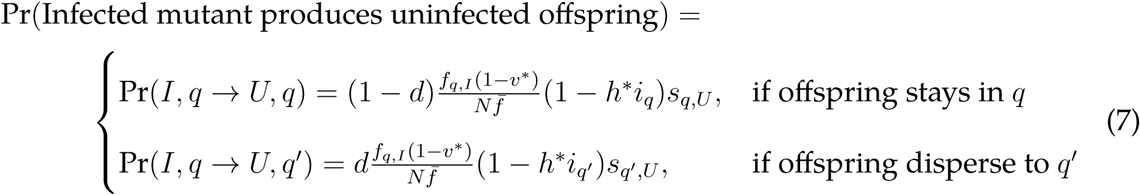

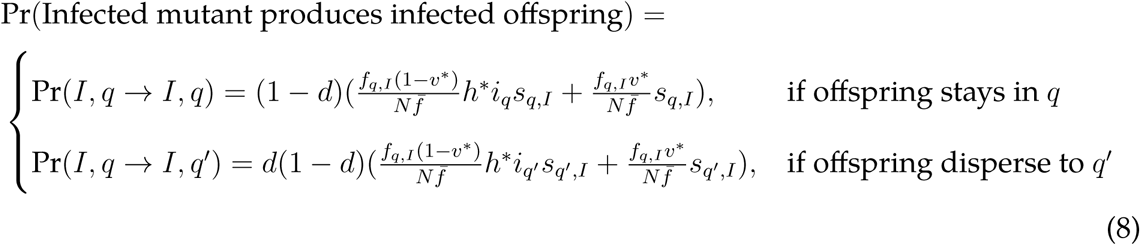

Using the above probabilities of mutant reproduction, we can write *A* as

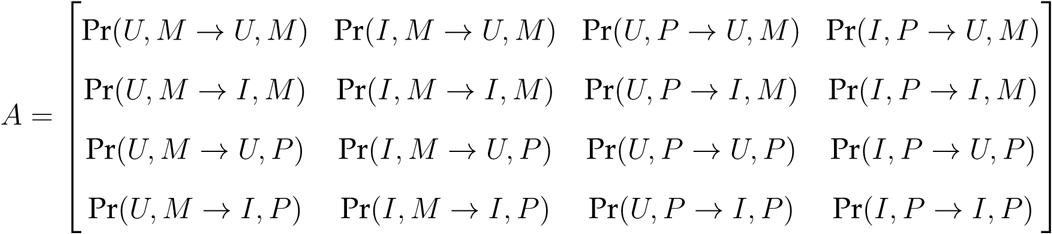

Because all cases in Equations 5 - 8 have a 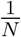 term, we can re-write *A* as

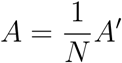

Unlike *A*, *A′* does not depend on *N*.

To find *B*, we start from the fact that, if a newborn establishes in patch *q*, an adult host in the patch has a 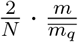 chance of dying (since there are 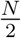 hosts in each of patch M and P). Because the population is comprised almost entirely of residents, the probability that a newborn establishes can be approximated using the probability that a newborn resident establishes. For patch *q*, where the other patch is *q′*, a host (mutant or resident) with mortality *m* has a probability of dying of

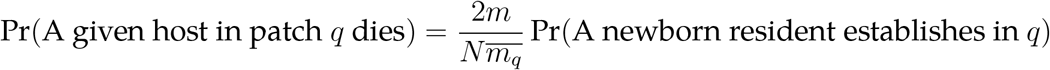

where

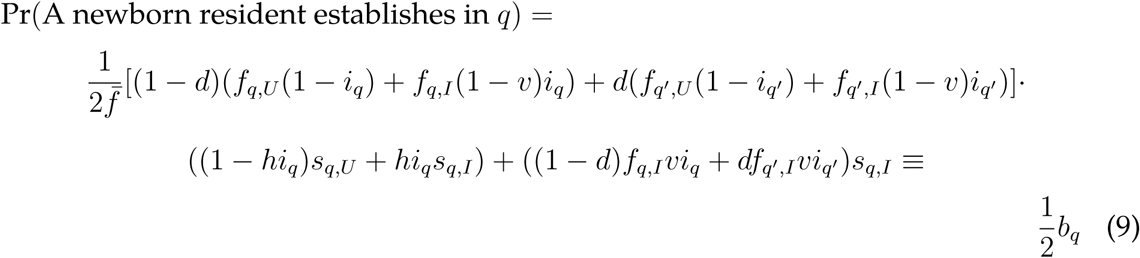

The 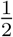 in the probability a resident establishes is due to the fact that each patch represents only half of the population and thus has its probability of reproducing normalized by 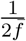. We separate it out from the rest of the expression (*b_q_*) to make it easier to deal with *A* - *B* later. This gives

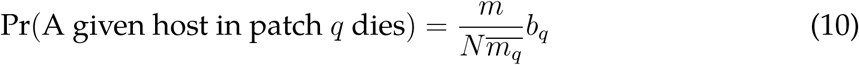

We can then write *B* as

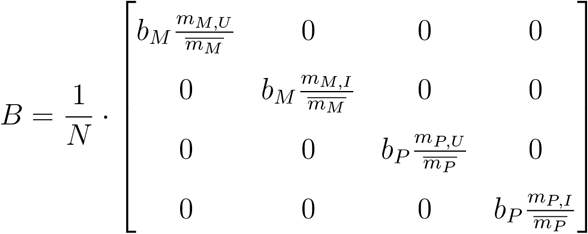

All the nonzero entries of *B* have a 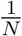 term. We can re-write *B* in terms of 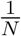 and *B′*, a matrix that does not depend on *N*.

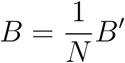

Then we can write *X_t_* as

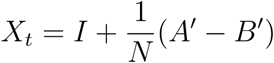

One problem with *X_t_* is that as *N* → ∞, *X_t_* → I. To fix this, we rescale time in units of *τ* = *tN*. Then the expected number of mutants produced per mutant of each patch and infection status can be written as

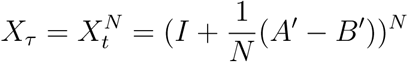

As the population size goes to infinity, we get the following formula for *X_τ_*

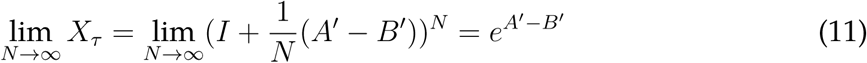

The mutant should invade if the leading eigenvalue of *X_τ_* > 1 when the resident is at equilibrium. Assuming mutations in transmission mode are small, we can trace the evolutionary trajectory of a population by seeing which mutant with similar transmission rates can invade, and then looking to see what transmission rates allow invasion of that mutant when it is the resident. Practically, this means finding the derivative of the leading eigenvalue of *X_τ_* at a range of resident transmission rates (a positive derivative means a mutant with a slightly higher transmission rate can invade, and a negative derivative means one with a lower transmission rate can invade). We then use these derivatives to trace the path of transmission mode evolution.

#### A.3 Transmission Mode Evolution - Symbiont Control

When transmission is a symbiont trait, we again investigate the invasion fitness of a mutant with slightly different transmission rates than the resident. We will follow the same general procedure as for host control. However, since a mutant symbiont should spread in the population if it can infected more hosts than the resident symbiont, we will track the number of mutants in units of hosts infected.

Let *X_t_* be the expected number of hosts infected with mutant symbionts in patches M and P by a mutant symbiont in each patch. The first and second rows of *X_t_* will give the infections produced in patches M and P, respectively. The columns of *X_t_* will likewise correspond to the location of the symbiont that produces the new infection.

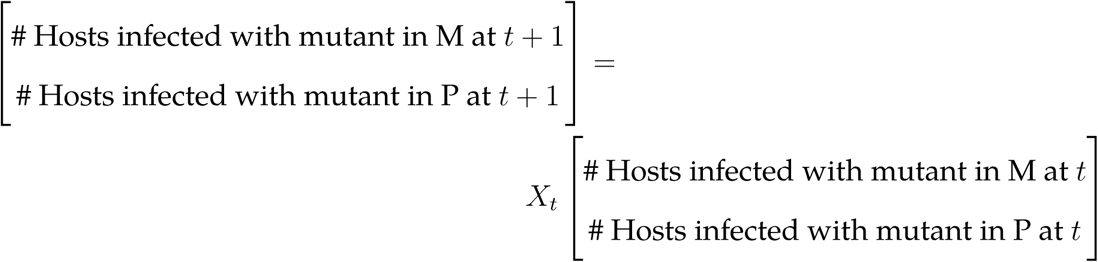

We can again define *X_t_* = *I* + *A* − B, where *A* is a matrix that gives the probability that a mutant symbiont produces a new in infection in each patch, and *B* gives the probability that a host infected with the mutant dies. Because a symbiont can produce an infection via horizontal or vertical transmission, we will write *A* as the sum of *A_υ_* and *A_h_*, the probability a mutant produces a new infection via vertical or horizontal transmission. We can get *A_υ_* from the probability a newborn host is born infected (Equation 1) and the probability a host born infected establishes (Equation 2).

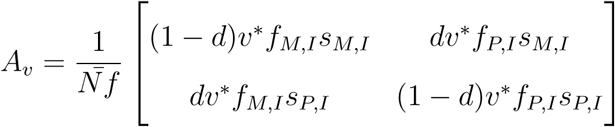

Because horizontal transmission is local, infections produced by horizontal transmission can only appear in the same patch as the original mutant symbiont, meaning *A_h_’*s off-diagonal entries will be 0. Infections produced by horizontal transmission depend both on the mutant’s horizontal transmission rate, its chance of being chosen as the newborn’s infectious contact (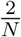), and on the number of incoming symbionts that are uninfected. The probability that a host is born uninfected in turn depends on the resident’s vertical transmission rate (*υ*). The diagonal entries of *A_h_* will then be

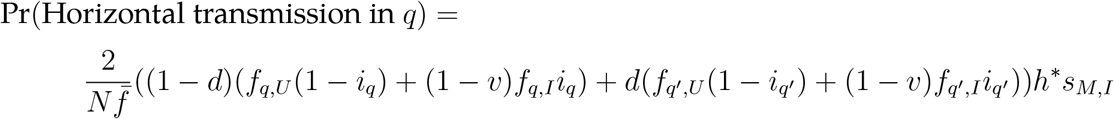

where *q* is the patch the host is arriving in and *q′* is the other patch. Then,

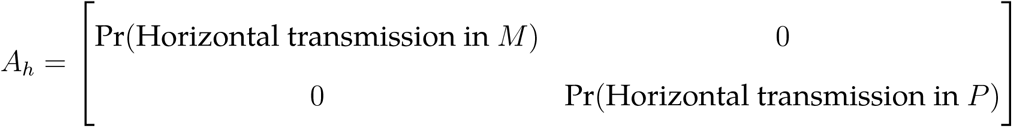

The probability that a mutant symbiont dies depends on the rate of newborn hosts establishing in its patch. This is given by Equation 10, which will be the diagonal entries of *B*. (As in the host case, the off-diagonal entries of *B* will be 0.)

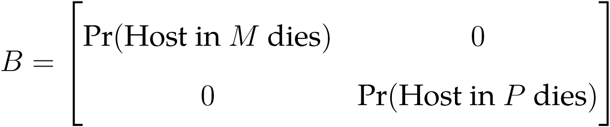

We can now see that *A* = *A_υ_* + *A_h_* and *B* have 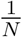 terms in them. We can re-write *A* and *B* as 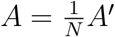 and 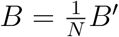, where *A′* and *B′* do not depend on *N*. Then the growth rate of a mutant symbiont in time units of *τ* = *tN* is *X_τ_* = *e^A′−B′^* as *N* → ∞.

Again the mutant should invade if the leading eigenvalue of *X_τ_* > 1 when the resident is at ecological equilibrium.

